# Calcimycocavitological Studies on Seashells from Beaches of North Goa, India

**DOI:** 10.1101/215228

**Authors:** Sujata Dabolkar, Nandkumar Kamat

## Abstract

Calcimycocavitology deals with the study of hollowing out of spaces in hard calcareous seashells by the fungi called calcimycocavites. Endolithic fungi in the shells were first reported and named as the trace fossils in 1889 by Bornet and Flahault. Endolithic fungi bore inside the shell by the process of bioturbation by using organic acids (oxalic acid, citrate) and enzymes (proteases, dehydrogenases and oxidoreductases). Previous reports show the presence of tunnels formed by fungal hyphae in the shells. The present work reports preliminary results of calcimycocavitological studies on the seashells from beaches of north Goa, India. The calcareous sand samples were collected from Arambol, Ashvem, Morjim, Vagator, Anjuna, Baga and Miramar beaches by pool sampling method and were separated into different fractions by using standard sieves. Each fraction of sand was subjected to stereomicroscopic studies which revealed that the sand fraction between 150-250 μm included irregular shell fragments showing positive colonization by calcimycocavites. Hydrochloric acid treatment was used to dissolve the shells and release calcimycocavites biomass which was stained with Congo Red and tentatively identified as distinct microfungal forms. SEM studies of the calcareous shell fragments revealed the microtunneling behavior of the fungi. Digital analysis of SEM images using Mountain premium 7.2 software revealed the fine topography of calcimycocavites hyphae along with unidentified presumptive biomineral encrustations. The ecological, biological and biogeochemical implications of the findings are presented with respect to possible role of calcimycocavites in Calcium and Carbon cycling by breakdown of the calcareous shells and release of inorganic and organic components in the ecosystem.

## INTRODUCTION

This work presents preliminary results of surveys undertaken from May 2014 to September 2015 aimed at production of a pictorial atlas of calcimycocavites from beaches of Goa and extend the knowledge to other areas in India and Asia. Sea shells are made up of organic matrix consisting of a proteins called conchiolin, a protein related to keratin (Hunt et al., 1978), glycan and biogenic calcium carbonate. Shells are the external skeletons of mollusks, an ancient and diverse phylum of invertebrates that was present in the earliest fossil record of multicellular life from the Cambrian period, over 500 million years ago. Diversity of sea shells of beaches in Goa beaches includes bivalves, Murex, Turis sumilis, mesogastropod shells, Turrittela sp. Fragmentation of sea shells occurs by physical, mechanical, chemical, biological processes such as by boring microorganisms, and by meteorological processes. Aim of this work was to carry out the Calcimycocavitological studies on the seashells which deals with the hollowing out of spaces in hard calcareous seashells by the fungi called calcimycocavites. Literature has shown the appearance of chytrids fungus such as *Saccomorpha clava* and *Polyactina araneola* in the sea shells of Australian beach sand (Zebrowski et al., 1937). The fungus *Neurospora crassa* has been found to be involved in the process of biomineralization of metal carbonates by using enzymes such as urease (Li et al., 2014). Every year a very heavy debris deposition is seen on beaches of Goa specially in calcareous sand. As shown in the table 1 there are less studies on the Endolithic fungi all over the world. During this work efforts were made to collect, classify and carry out microscopic studies of the seashells. Hydrochloric acid treatment was used to dissolve the shells and release calcimycocavites biomass which was stained with Congo Red and tentatively identified. SEM studies of the calcareous shell fragments and digital analysis of SEM images using Mountain premium 7.2 software were carried out. This work also helped in knowing the important role of calcimycocavites in calcium and carbon cycling. This may have wide application in ecology, biology and biogeochemistry.

**Table 1:** Previous work on Endolithic fungi

| Important findings | References |
| --- | --- |
| Lime-loving molds from Australian sands | Zebrowski et al., 1937 |
| The role of marine fungi in the penetration of calcareous substances | Kohlmeyer, 1969 |
| Fungi in corals, black bands and density-banding of *Porites lutea* and *P. Lobata* skeleton | Priess et al., 2000 |
| Endolithic fungi from deep sea calcareous substrata: isolation and laboratory studies | Raghukumar et al., 2002 |
| Endolithic fungi in marine ecosystem | Golubic et al., 2005 |
| Why are some microorganisms boring | Cockell et al., 2007 |
| Microborings from shallow marine habitats on both sides of the Panama Isthmus | Radtke et al., 2010 |
| Survey of shell-boring microorganisms Across a depth gradient at point caution, on San Juan Island, WA | Johnson et al., 2011 |
| Biomineralization of metal carbonates by *Neurospora crassa* | Li et al., 2014 |

## MATERIALS AND METHODS

Goa is the second smallest state located on the west coast of India covering an area of 3702 sq kms and runs 105 km long and 65 km wide. Goa is located between latitudes 14º 53’ 55’’N and 15º 47’ 59’’N and longitudes 73º 40’ 34’’E and 74º 17’ 03’’E. The Arabian Sea marks the western boundary of the State. Survey, exploration and selection of suitable sampling sites with heavy sea shell debris deposits was carried out using Google earth (Fig 1). Sampling of the sand with heavy seashell debris by pool sampling method. Separation of sea sand into different fractions was carried out by using standard sieves. The selection of seashell fraction with presumptive calcimycocavites was carried out. The selected shell fragments were mounted with DPX and visualized for the presence of calcimycocavites. Scanning electron microscopy(SEM) studies of calcimycocavites was carried out by using Golubic et al., 2005 method. Endolithic colonization was studied using dissolution by Hydrochloric acid method (Kohlmeyer, 1969) and microscopic studies. Digital Image analysis of SEM images was carried out using Mountains Premium 7.2, which helped in study of 3-D function, Pseudo images and also in finding the length and breath. 24bitmapped images were processed using SCION software (4.0.2) for following parameters. 1. Find edge function output, 2. The density slice function output, 3. The histogram profile (HP) and 4. The surface pixel plot density (SPPD).

**Figure 1:**
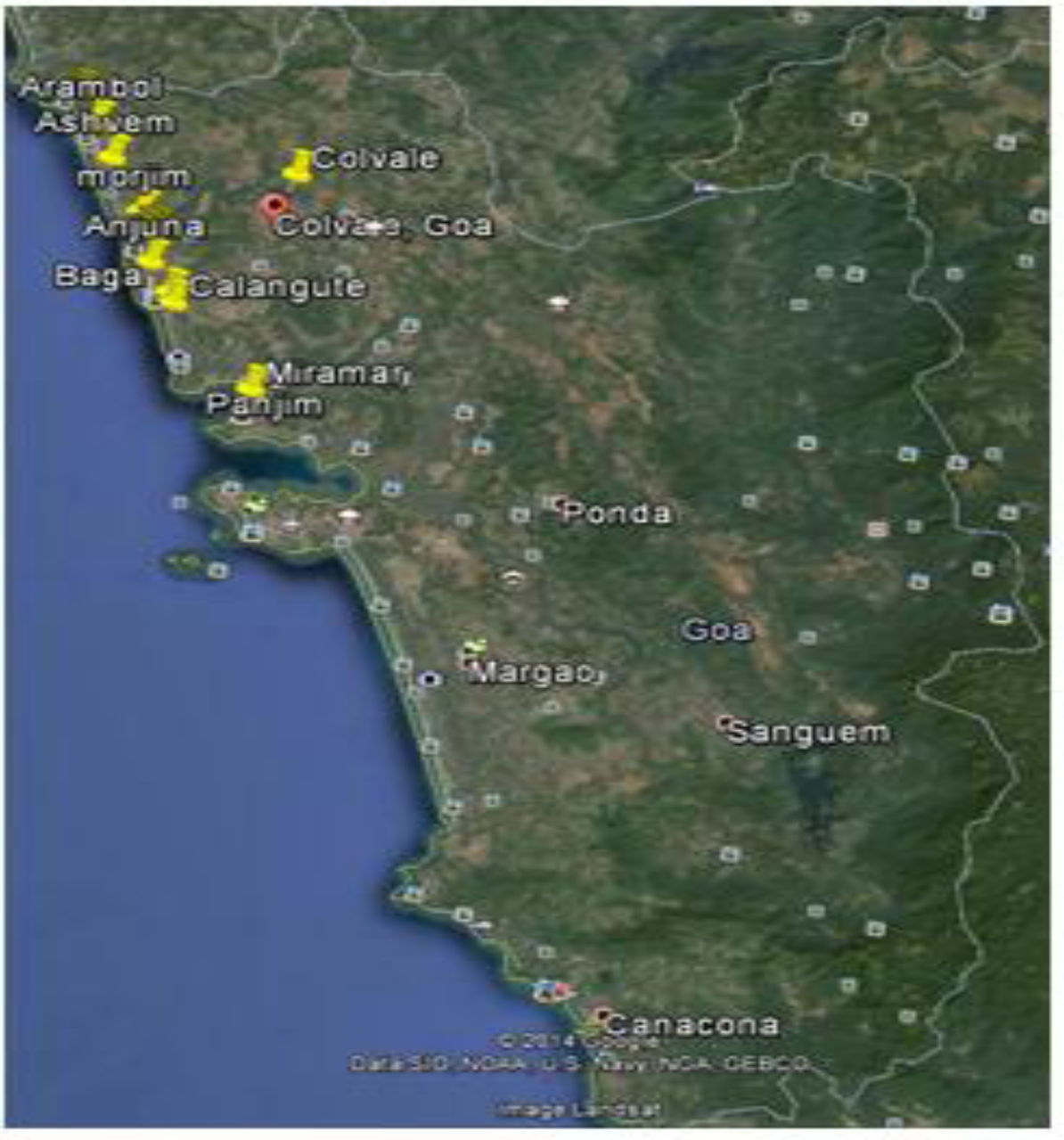
Location of shell fragment sampling sites

## RESULTS AND DISCUSSION

In all seven beach samples from North Goa yielded very small fractions containing infected fragments. On random examination of 1000 samples, 250 infected fragments were studied in detail to reveal the presence of calcimycocavites. Calcareous shell fragment showed the presence of calcimycocavites which were involved in colonization and formation of micro tunnels inside the shells (fig 2a-2d). Unidentified Endolithic fungi showed the presence of exopolysaccharide stained strongly in Congo red (Fig 3a-3d). Endolithic fungi showed the loose biomass (fig 3a-3c), compact matrix (fig 3b-3d) and strongly stained septate with monililoid branch (fig4a-4d). Output of Digital image processing revealed the typical septate mycocavite with presence of thick walls and thick septa (fig 5a-5g). Fig 6a-b shows the shell fragment topography, whereas 6c-6d shows the evidence of microbioturbation by calcimycocavite resulting in bores and tunnels as well as compact strands of thick walled hyphae of mycocavites (6e-6f). Fig 7a-Mycocavitological colony, 7b-Representative single hypha involved in mycotunneling. c–d: Hyphal mycometry. Fig7(e-g) and 8(a-g) shows output of digital image process.

**Figure 2(a–d):**
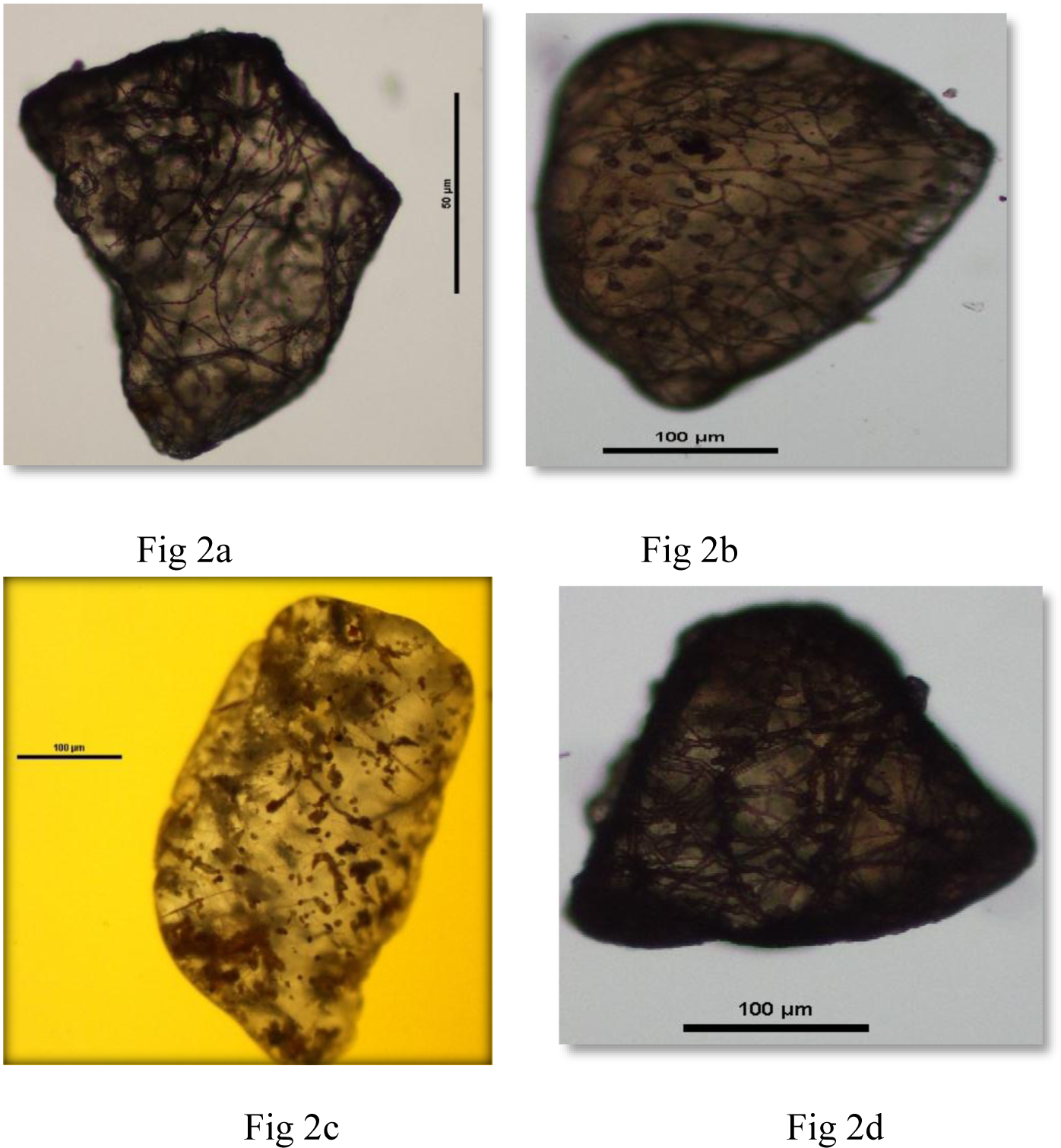
Calcareous shell fragments showing the presence of calcimycocavites, Colonization and microtunneling.

**Figure 3(a-d):**
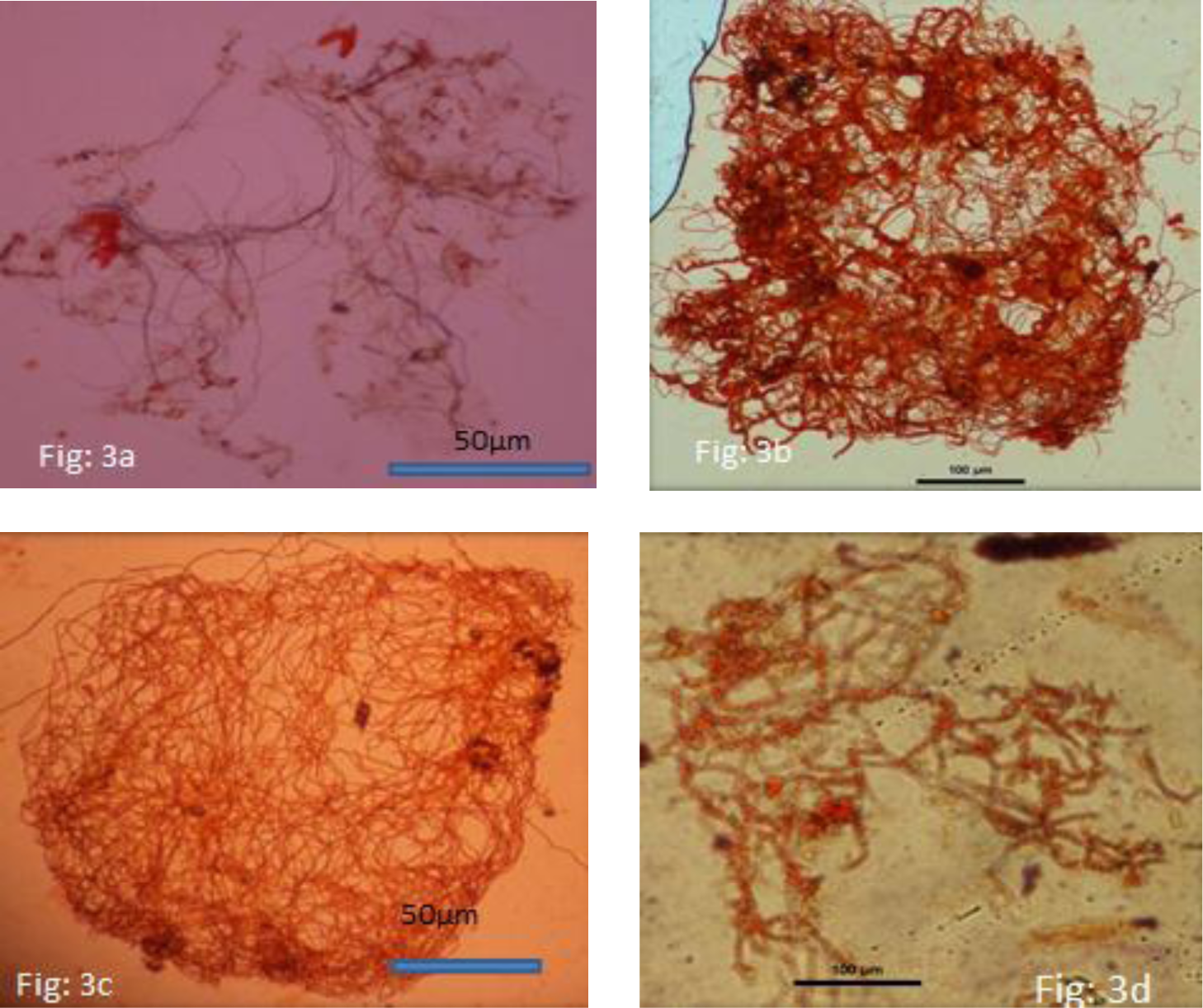
Unidentified endolithic fungi showing presence of exopolysacharide stained strongly in Congo red. a, c indicates loose biomass; b, d-indicates compact matrix.

**Figure 4(a-d):**
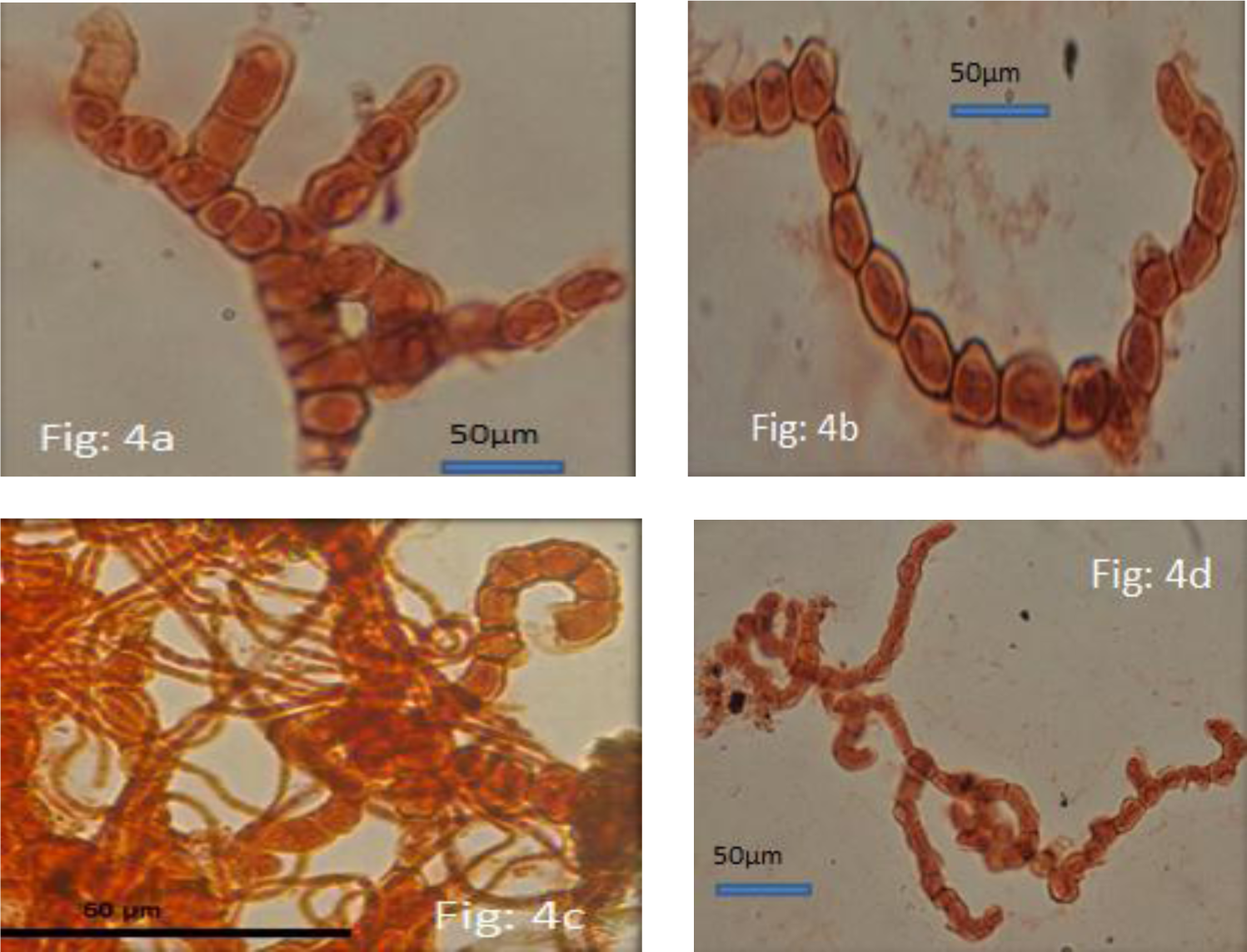
Strongly stained septate monililoid branched fungi

**Figure 5(a-g):**
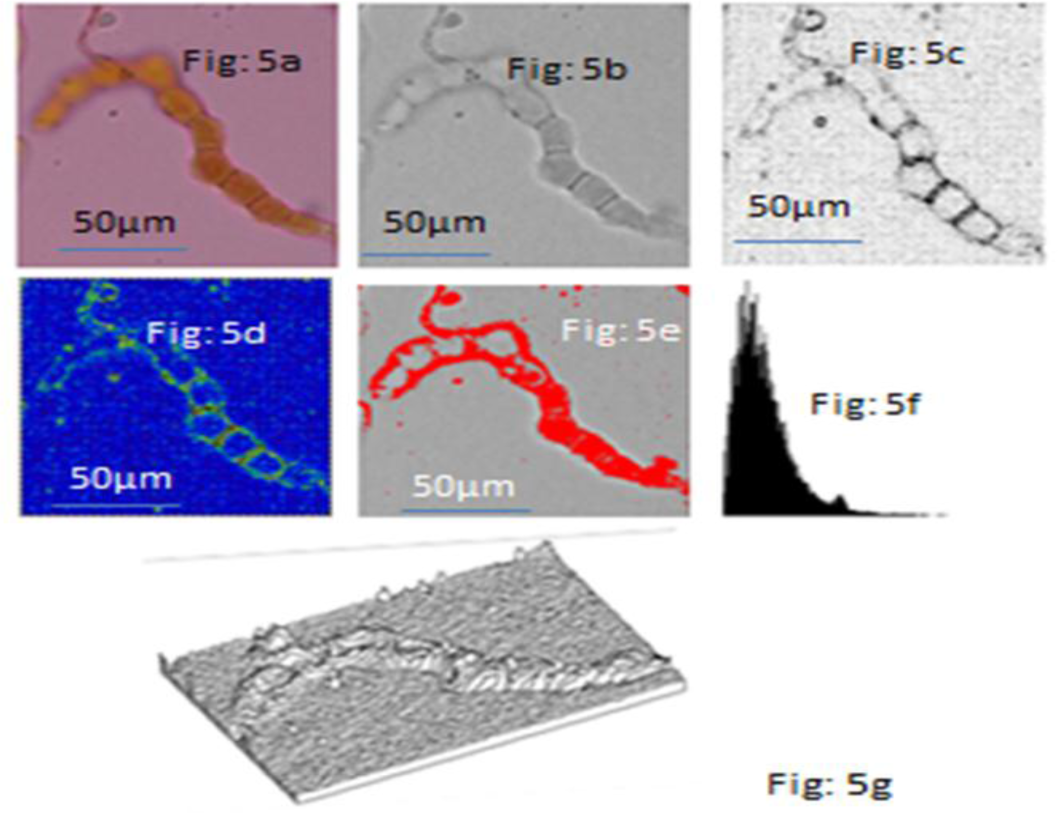
Output of SEM digital image processing of typical septate mycocavite with thick walls and thick septa. a-original hyphae, b-Negative, c-sharp edges of hypha, d-septation, e-density slice, f-histogram profile, g-surface plot.

**Figure 6(a-f):**
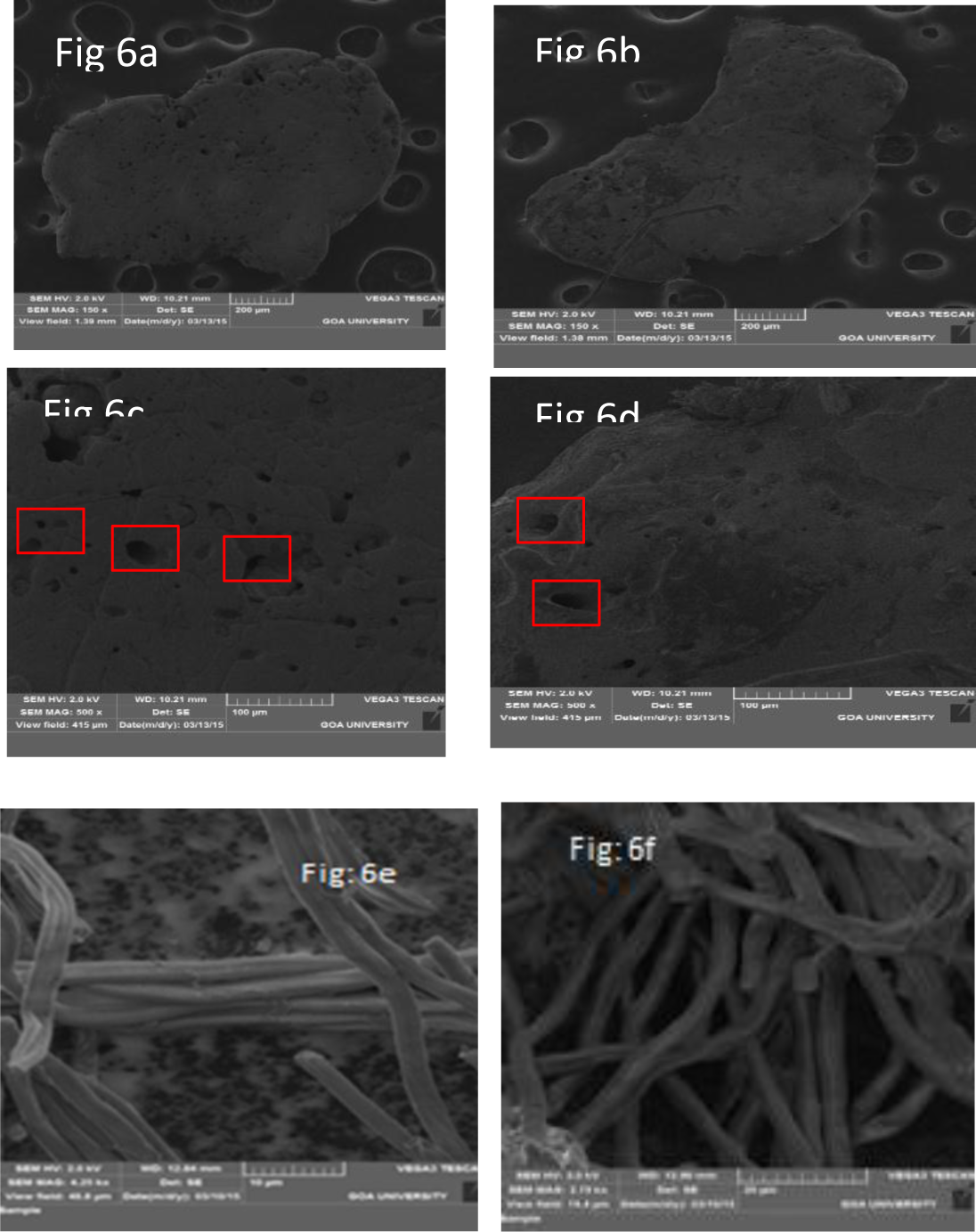
a,b-shell fragment SEM topography; c,d-Evidence of microbioturbation by calcimycocavite resulting in bores and tunnels; e,f-compact strands of thick walled hyphae of mycocavites.

**Figure 7(a-g):**
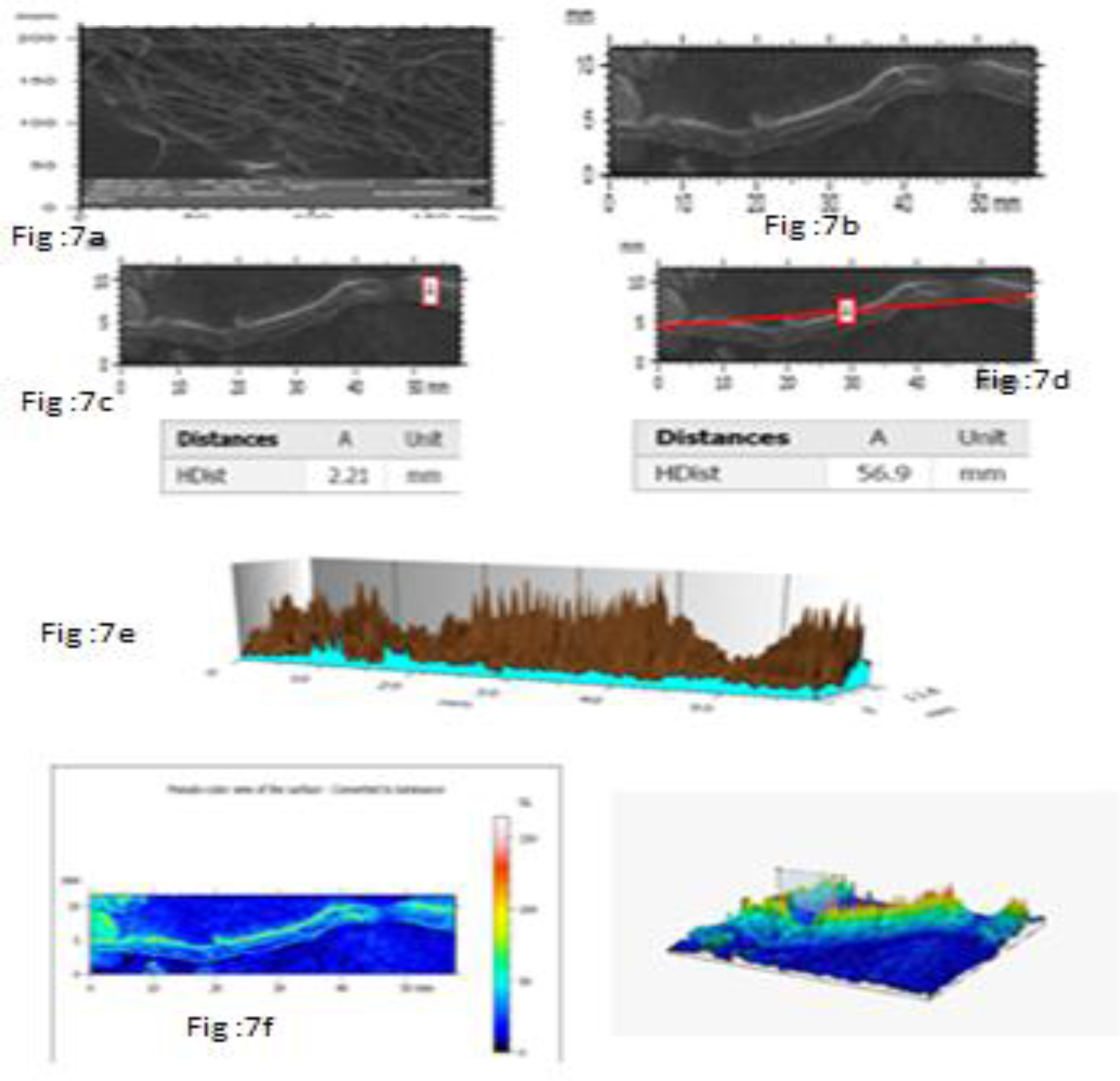
a-Mycocavite colony, b-Representative single hypha involved in microtunneling. c, d: Hyphal micrometry; e-g:Output of digital image process.

**Figure 8(a-g):**
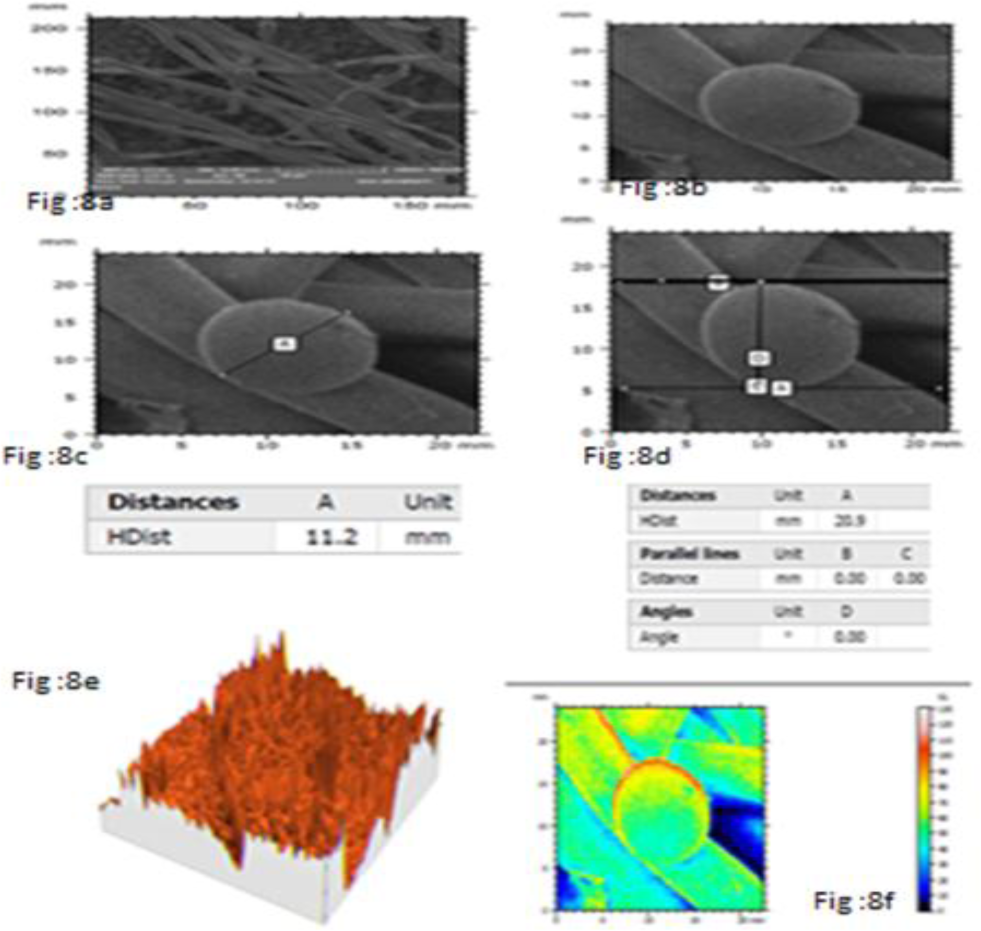
a-Mycocavite colony, b-Representative single branched initial. c, d: branched initial micrometry. e-g :Output of digital image process.

Seashell fragments on the beaches are infected by fungi probably due to mycelogenic units gaining entry by anchoring and penetrating through eroded regions in hyperhaline environment and spend indeterminate time as slow growing, chemoorganotrophic endolithic colonies (along with other unidentified members of microbial community such chemolithotrophic, archea, eubacteria and actinobacteria), using organic acids like Oxalic acid (Bech-Andersen, 1987; Greenet al., 1991; Larsen and Green, 1992; Jarosz-Wilkolazka and Gadd, 2003; Hastrup et al., 2006) to dissolve the Calcium Carbonate shell biomineral matrix to create microtunnels/boring, while depositing encrustrations of insoluble Calcium Oxalate. For growth and nutrition these mycocavites secrete extracellular hydrolytic enzymes to utilize shell proteins and glycan’s as source of nutrients. Microtunneling process mediated by dissolution of Calcium Carbonate causes the fragments to become fragile, disintegrate leading to the dispersal of the vegetative cells and asexual spores of calcimycocavites. These can again infect other fragments. Unless removed by tidal action flushing the sea fragments, the biodegradation of mycocavitogenic insoluble Calcium Oxalate by oxalotrophic members of bacterial communities may occur using bacterial Oxalate decarboxylase and formate dehydrogenase finally releasing Carbon Di Oxide and water and Calcium oxide to local nutrient pool. Calcimycocavites may be thus involved in biogeochemical cycling of Carbon and Calcium in Ocean-Beach environment. A scheme speculating the role of calcimycocavites in a typical beach ecosystem (Dabolkar and Kamat, 2015) has been explained (Fig 9).

**Figure 9:**
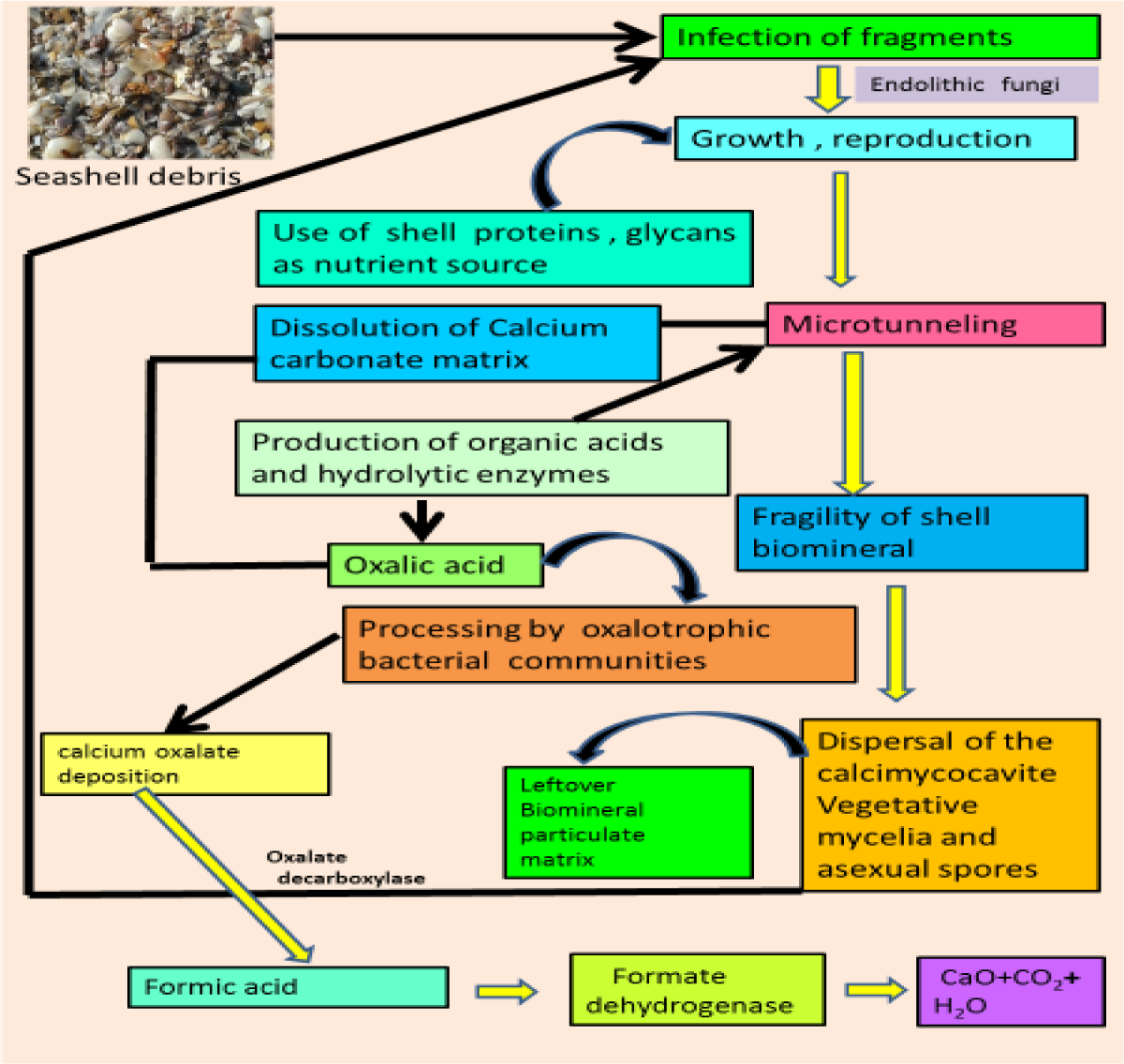
The postulated role of calcimycocavites in a typical beach ecosystem

## CONCLUSIONS

The results show that using combination of simple field and laboratory techniques and advanced software it is possible to visualize the endolithic fungi and undertake microbioturbation studies on shell fragments which are difficult to date. It has to be seen whether such calcimycocavites are distinctly related to marine fungi or unique to beach mycoflora but definitely they have an interesting source. Attempts are in progress to recover the calcimycocavites in viable form using novel media and cultural techniques so molecular methods can be used for their identification. Their role in biogeochemical cycling of Carbon and Calcium through Carbonate-Oxalate cycle is implied (while not denying the role of hitherto unidentified associated members of endolithic and epilithic microbial community). This adds a little weight to present discourse in global warming, climate change and role of fungi in Carbon and Calcium cycling at Ocean-Land interface. It has not escaped our notice that several of these microfungi could be trace microfossils, interesting from paleonotological viewpoint, perhaps helping to shed light on paleobiodiversity of fungi and paleoenvironment (Golubic et al., 2005). Explosive release of entrapped gas bubbles is another phenomenon from colonised shell matrix indicating their ecological and biogeochemical importance. Micro sampling of these bubbles using techniques used in sampling trapped bubbles in drilled cores of ice may yield valuable data on paleoclimates. We are in the process of refining the technique for using the calcimycocavites as biomarker proxies for different tropical beach environments. Such proxies may help in efficient environmental biomonitoring of the beach ecosystems.

## ACKNOWLEDGEMENTS

We thank Anne Berger, Sales Manager,Digital Surf, France for giving permission to use Mountains Map software, Digital surf for SEM image processing and analysis. This work was supported by UGC-SAP Phase II – Biodiversity, Bioprospecting programme and Goa University Fungus Culture Collection (GUFCC). We thank R.N.S Bandekar CO, Vasco da Gama for funding the work on biomineral studies.

